# Channelrhodopsin C1C2: Photocycle kinetics and interactions near the central gate

**DOI:** 10.1101/807909

**Authors:** M. R. VanGordon, L. A. Prignano, R. E. Dempski, S. W. Rick, S. B. Rempe

## Abstract

Channelrhodopsins (ChR) are cation channels that can be expressed heterologously in various living tissues, including cardiac and neuronal cells. To tune spatial and temporal control of potentials across ChR-enriched cell membranes, it is essential to understand how pore hydration impacts the ChR photocycle kinetics. Here, we measure channel opening and closing rates of channelrhodopsin chimera and selected variants (C1C2 wild type, C1C2-N297D, C1C2-N297V, and C1C2-V125L) and correlate them with changes in chemical interactions among functionally important residues in both closed and open states. Kinetic results substantiate that replacement of helices I and II in ChR2 with corresponding residues from ChR1, to make the chimera C1C2, affects the kinetics of channelrhodopsin pore gating significantly, making C1C2 a unique channel. As a prerequisite for studies of ion transport, detailed understanding of the water pathway within a ChR channel is important. Our atomistic simulations confirm that opening of the channel and initial hydration of the previously dry gating regions between helices I, II, III, and VII of the channel occurs with 1) the presence of 13-*cis* retinal; 2) deprotonation of a glutamic acid gating residue, E129; and 3) subsequent weakening of the central gate hydrogen bond between the same glutamic acid E129 and asparagine N282 in the central region of the pore. Also, an aspartate (D292) is the unambiguous primary proton acceptor for the retinal Schiff base in the hydrated channel.

**SIGNIFICANCE:** Channelrhodopsins (ChR) are light-sensitive ion channels used in optogenetics, a technique that applies light to selectively and non-invasively control cells (e.g., neurons) that have been modified genetically to express those channels. Using electrophysiology, we measured the opening and closing rates of a ChR chimera, and several variants, and correlated those rates with changes in chemical interactions determined from atomistic simulations. Significant new insights include correlation of single-point-mutations with four factors associated with pore hydration and cation conductance. Additionally, our work unambiguously identifies the primary proton acceptor for the retinal chromophore in the channel open state. These new insights add to mechanistic understanding of light-gated membrane transport and should facilitate future efforts to control membrane potentials spatially and temporally in optogenetics.

## I. INTRODUCTION

Channelrhodopsins (ChRs) mediate the first committed step in phototactic behavior, the mechanism by which unicellular algae migrate toward optimal growth conditions to transform light energy into complex substances such as amino acids and carbohydrates. ChRs aggregate in an eyespot region of the plasma membrane of algae, such as *C. reinhardtii*, and conduct monovalent and divalent cations down their electrochemical gradients upon light activation. While phototaxis has been investigated for over one hundred years, molecular identification of channelrhodopsins has occurred only recently. That molecular data strengthened the proteins’ prominence in the field of optogenetics, a biological technique that uses light to control cells in living tissues.^1,2^

Experiments have shown that channelrhodopsins can be expressed heterologously in various cell types, including cardiac and neuronal cells, thus enabling spatial control of cardiac muscle contraction and action potentials. ^3,4^ Temporal regulation of the ChR-enriched excitable cell membranes depends on opening and closing rates of ChRs. Channelrhodopsin photocycle kinetics have been described previously by a four-state kinetic model with two closed and two open states (Fig. 1).^5^ Upon extended time in the dark state, channelrhodopsin accumulates in the closed state, C1. Application of light to ChRs causes all-*trans* to 13-*cis* retinal isomerization and yields a high-conducting open state, O1. Upon extended light activation, equilibrium is reached between O1 and a lower conducting state, O2. Once light is turned off, the ion channel closes and transitions to the C2 state, and eventually converts back to the initial C1 ground state.

**Figure 1.**
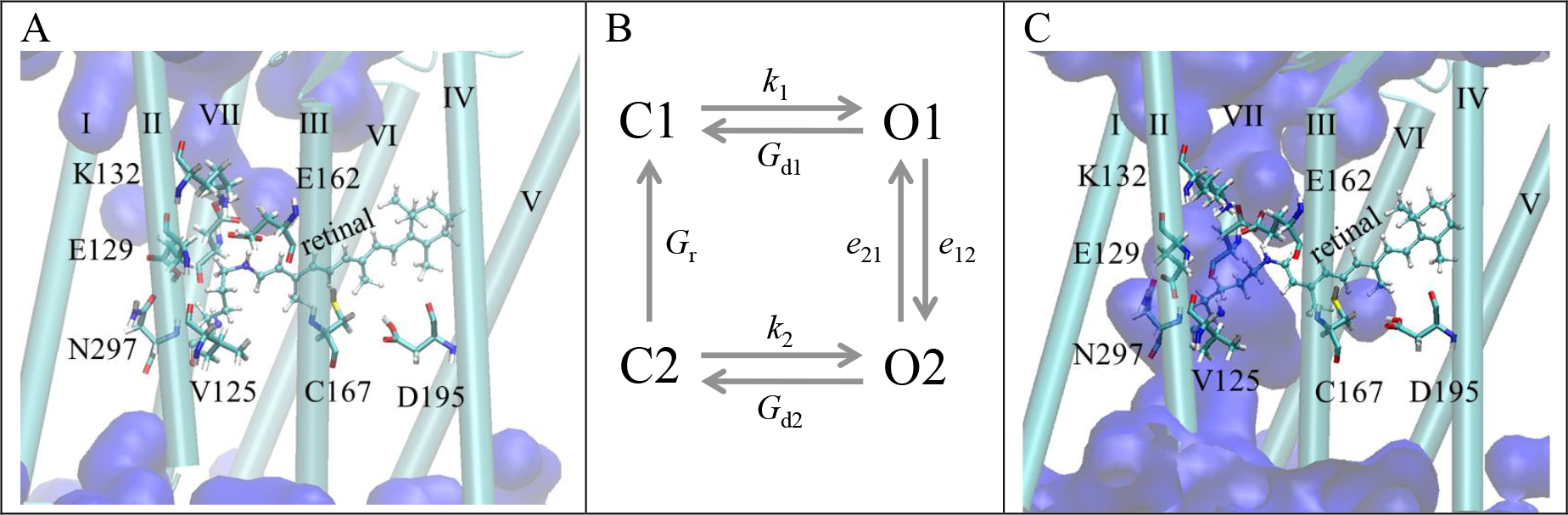
A) Simulation snapshot of closed-state wild type channelrhodopsin chimera (C1C2-WT). B) Four-state ChR2 photocycle model, with two closed (C1 and C2) and two open (O1 and O2) states. *G*_d1_ and *G*_d2_ are the rates of O1→C1 and O2→C2 transitions, respectively. *G*_r_ is the rate of thermal C2→C1 conversion and recovery of the initial closed state. The rates of transitions between open states 1 and 2 are *e*_12_ and *e*_21_. Activation rates *k*_1_ for C1→O1 and *k*_2_ for C2→O2 processes are associated with protein conformational changes related to absorption of a photon of light. C) Simulation snapshot of open-state C1C2-WT. Water, represented as blue surfaces, fills the open channel, but not the closed channel.

The first crystallized light-activated ion channel consisted of a dark-adapted, non-conducting C1C2 chimera (Fig. 1A). This protein contains the first five transmembrane helices of channelrhodopsin-1 and the last two transmembrane helices of channelrhodopsin-2. Sharing a similar overall topology, both C1C2 and ChR2 comprise a putative pore among helices I-III and VII. In addition, mutagenesis and functional studies have demonstrated that both proteins accommodate numerous glutamic acid residues on helix II, which contribute to the ion selectivity, but have little influence on ChR2 photocycle kinetics.^6^ Prior to resolving a crystal structure of ChR2, ^7^ C1C2 chimera was used to model ChR2. The similarities between ChR2 and C1C2 secondary structures made the choice of the structural template plausible. However, the presence of helices I and II of channelrhodopsin-1 in the chimera resulted in non-negligible differentiation between C1C2 and ChR2.

Subtle structural differences introduced by single point mutations in helices I, II, III, and VII alter the kinetics of the ChR photocycle. For instance, ChR2-E123T (ChETA) exhibits faster channel closing (4 ms vs. 10 ms in wild type, ChR2-WT) concurrent with a moderate decrease in photocurrent magnitude, and strong desensitization. ^8^ The C128X (X=T, A, or S) mutation in ChR2 transforms the protein into step-function opsin at a cost of significant extension of the open-state lifetime (*τ*_off_ up to 30 minutes). ^9,10^ ChR2-T159C ^11^ and ChR2-L132C ^12^ display slower channel off-kinetics, with associated increase in photocurrent magnitudes. While ChR2-H134R exhibits enhanced currents during prolonged stimulation, it also displays a 2-fold slower channel closing rate, resulting in decreased temporal precision of the variant when compared with ChR2-WT. ^3,13^

Despite the experimentally observed effect of mutations on pore kinetics, little is known about the mutation-induced changes in chemical interactions among key gating residues. To understand better the correlations between intramolecular interactions and functional differences among ChR variants, we conducted computational and electrophysiology studies of wild type C1C2 (C1C2-WT), and mutants C1C2-N297D, C1C2-N297V, and C1C2-V125L (Fig. 2) in their closed and open states. Single point amino acid replacement sites have been chosen at, and in the vicinity of, the C1C2 central gate. The central gate is defined by the hydrogen bond between E129(90)…N297(258).^14,15^ (Numbers in parentheses correspond to the ChR2 amino acid numbering scheme, a shift by 39 units.) In closed-state C1C2, this gating interaction holds helices II and VII together in the central region of the pore, thus preventing continuous channel hydration and subsequent ion permeation. The choice of mutations was driven by the following questions. What is the influence of side chain acidity on gating interactions in C1C2-N297D compared with C1C2-WT? How is gating achieved in the absence of the E129(90)…N297(258) hydrogen bond (C1C2-N297V)? Knowing that V86 contributes to ChR2 recovery to the ground state,^16^ how does a conservative replacement of the analogous residue, which is spatially proximal to the central gate, affect chemical interactions in C1C2-V125L?

**Figure 2.**
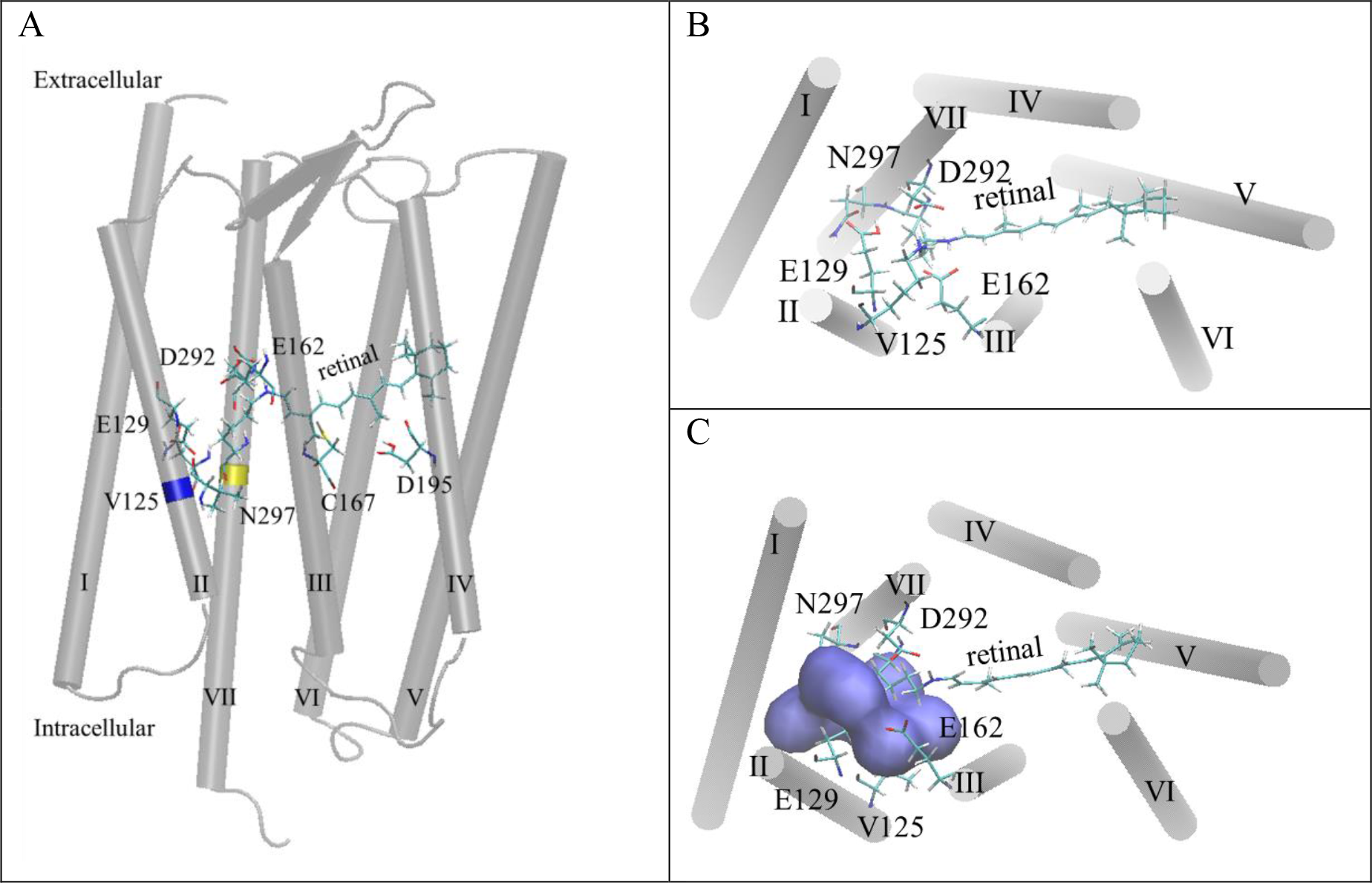
A) Location of mutated residues V125 (blue) and N297 (yellow) in the transmembrane region of equilibrated, closed-state C1C2-WT (gray cylinders). Key residues involved in protein gating are shown as sticks. For clarity, water, lipids, and ions are not shown. Simulation snapshot of equilibrated B) closed-state and C) open-state C1C2-WT seen from the extracellular side of the membrane. Water molecules present within 5 Å of the central gating residues, E129(90) and N297(258), are shown as blue surfaces. Residue locations: V125, E129 (helix II); E162 and C167 (helix III); D195 (helix IV); N297, D292, retinal (helix VII). Retinal is attached to helix VII through a covalent bond with K296 (helix VII), where a Schiff base forms.

## II. MATERIALS AND METHODS

### A. Molecular biology and expression of constructs in Xenopus oocytes

A plasmid containing the gene for C1C2 was obtained from Addgene ^17^(Addgene plasmid # 35519). The cDNA of the C1C2 chimera (residues 1-356) and C-terminal eYFP was subcloned into the vector pTLN. Xenopus oocytes were injected with 50 ng mRNA of wild type and mutant C1C2 for two-electrode voltage clamp experiments.^18–20^

### B. Data analysis and numerical fitting

Apparent kinetic parameters for channel activation and decay were determined by monoexponential fitting of photocurrent data from the time the light is turned on to peak current (*τ*_on_), and from peak to stationary current (*τ*_decay_) using Eq. 1(a). Fast (*τ*_fast_) and slow (*τ*_slow_) components of channel closing after the light is turned off were estimated by bi-exponential fitting from the steady-state to off current (*τ*_off_) using Eq. 1(b),

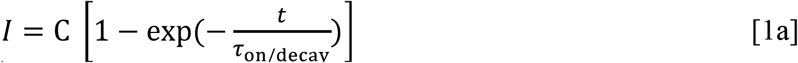

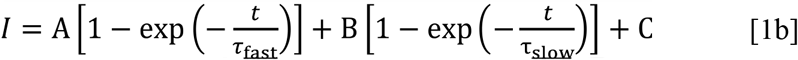

where *t* is the time in ms, *I* is the photocurrent in μA, *τ* is the time constant (in ms) of the associated transition, A, B, and C are constants.

The current-voltage relationship for each construct was determined from photocurrent traces recorded using the voltage step protocol (from membrane potential, *V*_m_, of −100 mV to +40 mV in +20 mV steps) in Na^+^ bath solution at pH 7.0 or pH 9.0 (115 mM NaCl, 2 mM BaCl_2_, 1 mM MgCl_2_ and 10 mM HEPES (pH 7.0) or 10 mM Tris (pH 9.0).

The reversal potential, *E*_rev_, is the membrane potential at which there is no net current flow through the channel. *E*_rev_ was found from the plot of photocurrent vs. membrane potential by linear interpolation, as previously described. ^19,16^ Data analysis was performed using Mathworks MATLAB R2017a.

### C. Four-state model fitting

Kinetic modeling of photocurrents was based on the four-state photocycle model (Fig. 1B). ^16,21^ Here, the model simulates channel photocurrents characterized by two closed states (C1 and C2) and two open states (O1 and O2) using the following set of rate equations,

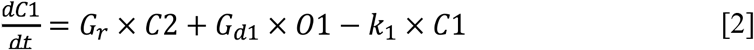

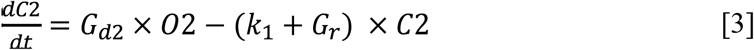

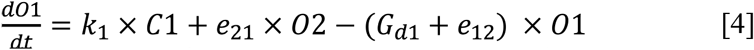

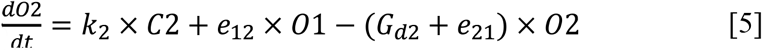

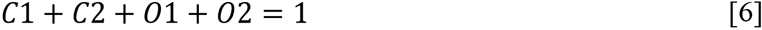

The variables *C1*, *C2*, *O1*, and *O2* represent the fractional population of each state. The rates of light-dependent channel activation given by *k*_1_ and *k*_2_ represent the transitions *C1* → *O1* and *C2* → *O2*, respectively. Rates for the reverse of these transitions are given by *G*_d1_ and *G*_d2_, respectively. The forward and reverse transitions between the two open states during prolonged light stimulation are represented by *e*_12_ and *e*_21_. Lastly, *G*_*r*_ represents the thermal conversion of *C2* back to *C1* in the dark. *Gr* was calculated by first quantifying the percent recovery of peak current (p) relative to stationary current (s): Δ*I*_2_/Δ*I*_1_ where Δ *I* _2_ = *I* _p2_ – *I* _s2_ and Δ *I*_1_ = *I*_p1_ – *I*_s1_. The subscripts correspond to the initial photocurrent (1) and current upon a second light pulse (2) following wait periods in the dark of 2, 4, 8, 16, and 32 seconds recorded at a holding potential of −100 mV. The percent recovery was plotted against the dark time interval to generate the recovery curve, and a monoexponential curve fit was performed to obtain the recovery time constant *G*_*r*_.^8^

### D. Simulations

Atomistic simulations were based on the crystal structure of the closed-state channelrhodopsin chimera, C1C2, with all-*trans* retinal (Protein Data Bank (PDB): 3UG9)^17^) protonated, E162(123) and D292(253) deprotonated. A molecular dynamics simulation box contained C1C2, lipid bilayer with 200 molecules (100 per layer) of 1,2-di-oleoyl-sn-glycero-3-phosphatidylcholine (DOPC), TIP3P water, and ions (Na^+^ and Cl^−^) to neutralize the protein net charge and provide 0.15 M salt concentration. The total number of atoms in the system was ~96,000. Electrostatic interactions were calculated using the particle mesh Ewald algorithm with a real-space cut-off distance of 12 Å and grid width of 1 Å. Single-point mutations V125L, N297V, or N297D were introduced to a previously equilibrated non-conducting C1C2-WT structure with all-*trans* retinal. All systems were equilibrated for 100 ns at 300 K. Protein structures and properties were calculated from statistical averages of molecular coordinates obtained from an additional 20 ns of the molecular dynamics (MD) production runs. We followed the previously reported molecular dynamics simulation set-up. ^22^ The p*K*_a_ values of the ionizable C1C2 chimera side chains were calculated with the PROPKA code and averaged over the last 20 ns of molecular dynamics simulations.

To generate the open state structure, a total of 13 C1C2-WT starting structures (Table S1) with 13-*cis* retinal, and varied protonation of E129(90), E162(123), D195(156), and D292(253) were tested to predict a configuration for an open pore. Varied protonation of the Schiff base (SB), a covalent link between retinal and lysine K296(257), and its orientation toward opposing sides of the pore were also tested. Deprotonation of D195(156) or the Schiff base (SB) yielded backbone root-mean-squared deviations (RMSD) of ~ 2 Å (Table S1) with respect to the pre-equilibrated structure. Continuous hydration of the channel was achieved by equilibration of C1C2-WT with 13-*cis* retinal, protonated Schiff base oriented toward the extracellular side of the channel (SBH) and E129(90), E162(123), and D292(253) deprotonated. The latter system’s backbone RMSD of 1.1 Å (Table S1, structure 13) was the lowest of all tested open-state C1C2-WT protonation state variants, and helix VII displacement upon pore hydration was confirmed with normal mode analysis (see Supporting Material, Fig. S1, for details). This result is in agreement with the E90-helix II- tilt mechanism in ChR2 reported by Kuhne.^14^ That structure was used for the open-state C1C2 studies carried out here. Differences in structure between the closed and open states were in the conformation of retinal and protonation of the gating residue E129(90) (Table S1).

## III. RESULTS

### A. Electrophysiological recordings

Functional expression of wild type and mutant C1C2 constructs in *Xenopus laevis* oocytes was assessed using a classic two-electrode voltage clamp. Each construct yielded measurable photocurrents (Fig. 3) upon activation with blue light (*λ*_max_ = 470 nm). Apparent kinetic rates were extracted from photocurrent traces (Fig. 3A) measured at a membrane potential of −100 mV by fitting either mono- or bi-exponential equations (Eq. 1) to the experimental data (Table 1). The recovery time constant (*G*_r_) of the transient peak current upon subsequent light pulses is important since a rapid recovery time is required to achieve fast repetitive stimulation of neurons. Recovery of the transient peak current is significantly slower for C1C2 (*G*_r_ = 0.53 ± 0.03 ms^−1^) than for ChR2 (*G*_r_ = 6.7 ± 0.1 s^−1^). ChR2 peak current recovers fully to its initial intensity within 30 s in the dark, whereas C1C2 plateaus at a maximum of 80% recovery. This result is consistent with partial initial maximum responses reported by Lin et al.^23^

**Figure 3.**
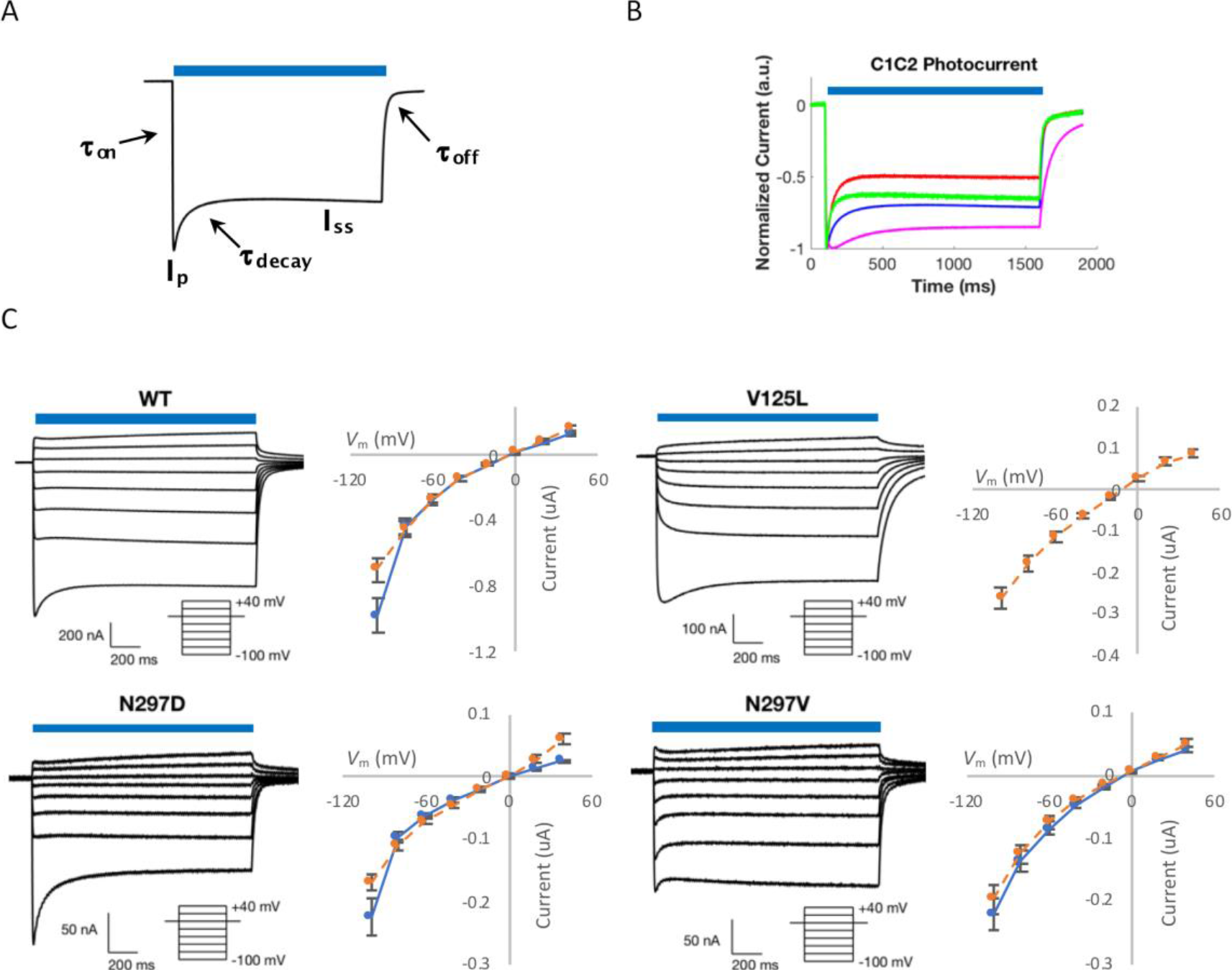
Kinetic analysis of photocurrents from two-electrode voltage clamp recordings. Light activation is indicated by the blue bars. A) Indication of characteristic parts of the photocurrent curve used to analyze channel kinetics. B) Normalized photocurrent traces of C1C2 constructs expressed in Xenopus oocytes, *V*_m_ = −100 mV, Na^+^ solution, pH 7.0. Light activation was achieved with a fiber-optic guided 470-nm LED light. C) Photocurrent traces for C1C2 wild type and single mutant constructs using the inset voltage protocol for Na^+^ measuring solution, pH 7.0. Scale bar is for each representative trace and current (*I*) - voltage (*V*) relationship for *I*_P_ (blue) and *I*_SS_ (orange) representing average ± standard error of the mean.

**Table 1.** Apparent kinetic rates, current ratios, and reversal potentials extracted from photocurrent traces recorded at *V*_m_ = −100 mV and Na^+^ measuring solution (pH 7.0). #

| | $\tau_{\text{on}}$ [ms] | Time<br>to<br>peak<br>[ms] | $\tau_{\text{decay}}$<br>[ms] | $\tau_{\text{off}}$<br>Fast<br>[ms] | $\tau_{\text{off}}$<br>Slow<br>[ms] | $G_r^\dagger$<br>[$\text{s}^{-1}$] | $E_{\text{rev}}$ (mV)<br>$I_p$ $I_{ss}$ | | $I_{ss}/I_p$ |
| --- | --- | --- | --- | --- | --- | --- | --- | --- | --- |
| C1C2-WT | $2.34 \pm 0.09$ | $16.7 \pm 1.1$ | $31.5 \pm 2.5$ | $11.2 \pm 0.5$ | $38.5 \pm 1.1$ | $0.53 \pm 0.03$ | -3.5<br>$\pm 0.9$ | $-4.1 \pm 0.7$ | $0.77 \pm 0.02$ |
| V125L | $2.9 \pm 0.2^*$ | $27.1 \pm 2.5^{**}$ | $89.0 \pm 8.4^{**}$ | $41.5 \pm 3.1^{**}$ | $131.9 \pm 8.3^{**}$ | > 30 min | ND | $-9.8 \pm 1.1^{**}$ | $0.90 \pm 0.02^{**}$ |
| N297D | $2.02 \pm 0.05^{**}$ | $11.7 \pm 0.4^{**}$ | $38.2 \pm 3.1$ | $9.3 \pm 0.3^{**}$ | $39.5 \pm 1.5$ | $0.6 \pm 0.2$ | -2.9<br>$\pm 0.6$ | $-2.3 \pm 1.0$ | $0.65 \pm 0.04^*$ |
| N297V | $2.5 \pm 0.2$ | $13.4 \pm 1.6$ | $53.7 \pm 6.4$ | $9.7 \pm 0.7$ | $38.2 \pm 1.0$ | $0.86 \pm 0.04^{**}$ | -5.1<br>$\pm 0.7$ | $-6.8 \pm 1.2$ | $0.67 \pm 0.01^{**}$ |

<sup>#</sup> Each value is an average of 5-22 cells +/- standard error. Probability values: \*, $p < 0.05$ , \*\*, $p < 0.01$ .
<sup>†</sup> Values for $G_r$ were calculated in $\text{Na}^+$ measuring solution (pH 9.0) as values were too slow to measure at pH 7.0.

Application of the four-state kinetic algorithm to C1C2 experimental photocurrent traces (Table S2) reveal that the N297D mutant has no significant change in any kinetic parameter outside of a decrease in the maximal photocurrent. In contrast, the V125L mutant decreases the rate of transition between O1 and the adjacent conformation states C1 and O2, including a slower *τ*_on_, *G*_d1_, *e*_21_ and *e*_12_. This result demonstrates that, upon making the V125L mutation, entry and exit to/from O1 is slower. This slowing likely contributes to the exaggerated loss of *I*_P_ upon repeated light stimuli (Fig. 3B). In contrast, inspection of the kinetic algorithm for the N297V mutant demonstrates faster *G*_d1_, *e*_21_ and *e*_12_. Each of these faster components transitions C1C2 out of the O1 states and thus peak currents can be observed even upon consecutive voltage pulses, as seen in Fig. 3B.

Substitution of valine 125 with leucine in C1C2 resulted in significantly slower overall kinetics compared to the wild type variant. C1C2-V125L channel recovery was affected most severely among all tested constructs, resulting in complete abolishment of peak currents upon a second light pulse (Fig. 3B). No evidence of peak recovery was observed after the wait period of up to 30 minutes in the dark. Similar trends were reported for the analogous ChR2-V86L mutant. ^20^

The C1C2-N297D mutant showed slightly faster on and off rates compared to C1C2-WT, but a similar current decay rate. The analogous mutation in ChR2-N258D resulted in significantly slower decay (*τ*_decay_ = 25.83 ± 3.07 ms) and off kinetics (*τ*_off_ = 27.42 ± 0.96 ms) compared to ChR2-WT. ^24^

No significant differences in apparent kinetic rates were observed in the C1C2-N297V mutant compared to wild type. A slightly larger recovery time constant (0.86 ± 0.04 s^−1^) was observed in N297V compared with the wild type. This result contrasted with the analogous mutation in ChR2-N258V, which led to a complete cessation of measurable photocurrents despite a robust expression of protein. ^16^

The reversal potential, *E*_rev_, is the membrane potential at which the current flow through the channel reverses direction. Thus, *E*_rev_ indicates the ease of ion passage through the pore. Reversal potentials (Table 1) were calculated from both peak and steady state photocurrents (Fig. 3C). Compared to C1C2-WT, reversal potentials were observed to be slightly less negative for N297D and slightly more negative for the N297V mutant in Na^+^ at pH 7.0. Reversal potentials for V125L were calculated only for the steady state current due to the lack of recovery of peak current after the first light pulse. Compared to wild type, V125L exhibited slightly more negative *E*_rev_.

The ratio of steady state to peak current, *I*_ss_ / *I*_p_, provides a measure of the degree of channel inactivation (corresponding to *τ*_*decay*_) upon continuous illumination and gives insight into the relative conductance of the two open states (O1 and O2). Steady-state to peak current ratios were calculated for each construct (Table 1) from photocurrents recorded at −100 mV in Na^+^ solution at pH 7.0. The *I*_ss_ / *I*_p_ ratio for C1C2 wild type (0.77 ± 0.02) is higher than for ChR2 (0.36 ± 0.01), indicating that C1C2 undergoes less inactivation during prolonged light exposure. Comparison of C1C2 mutants against the wild type channel revealed that V125L had an even higher *I*_ss_ / *I*_p_ ratio, while significantly lower values were seen for both the N297D and N297V constructs.

Overall, the experimental results show that C1C2 and its mutants do not behave like ChR2 or its analogous variants except in the case of V125L. To gain insight about differences observed here between C1C2 mutants and the wild type channel, we carry out molecular dynamics simulations and analyze structure, hydration, and chemical interactions in closed and open channels.

### B. Molecular dynamics results

#### 1. Structure and hydration of closed-state C1C2-WT and mutants

Introduction of V125L, N297V, and N297D single-point mutations to closed-state, dark-adapted C1C2 yielded minimal structural changes in molecular dynamics simulations. For example, little distortion occurred to protein backbone atoms (RMSD ~ 1 Å, Table S3) when compared with the closed-state C1C2-WT structure. Furthermore, the separations among helices involved in gating, II and VII (Table S3), remained about the same in the closed states. In the extracellular, central, and intracellular parts of the protein pore, those separations measured ~15, ~10, and ~10 Å, respectively. Finally, the central gate appeared mostly the same in two mutants (V125L and N297D) as in the wild type channel (Table S4). Recall that the central gate consists of a hydrogen bond between the side chains of protonated E129(90) on helix II and asparagine or aspartic acid N/D297(258) on helix VII. The N297V mutant was designed to eliminate that central gate hydrogen bond.

Similar patterns of partial hydration were observed among the closed-state C1C2 variants studied here. Notably, a water discontinuity existed consistently in the central region of the channel, near the gating residues E129 and 297. Also, partial pore hydration in all closed-state C1C2 variants was observed, with an average of 30-37 waters (Table S3) in the transmembrane region of the channels. Residue 125 is spatially proximal to the gating region of C1C2. In the closed-state channels, separations between helices II and VII in the V125L mutant and WT differed by only 0.3 Å or less in either extracellular, central, or intracellular sides of the pore. Nevertheless, C1C2-V125L accommodated on average 5 more waters compared with the WT channel in the closed states.

Given the similarity in structure and hydration, we attribute the experimentally observed differences in the photocycle kinetics among the studied C1C2 variants to the mutation-induced rearrangement in the network of interactions among functionally important residues.

#### 2. Interaction networks in closed-state C1C2-WT and mutants

Although the separations between the gating helices (II and VII) remained nearly the same among all the closed state channel structures, the mode of interaction changed significantly. As noted above, placement of valine in position 297(258) on helix VII prevented formation of the central gate hydrogen bond between residues 297 (helix VII) and 129 (helix II) (Table 2). Instead, E129(90) formed a hydrogen bond directly with E162(123) at a separation of *r*(O…O) = 2.6 ± 0.1 Å (Table S4). As a result of strong competition in hydrogen bonding between E162(123) and residues E129(90), K296(257), D292(253), T166(127), and K132(93), the distribution of acid dissociation constant values (p*K*_a_) for residue E162 showed a multimodal pattern, with peaks at 1, 4.5, and 10 p*K*_a_ units (Fig. S2). Further, the E162 p*K*_a_ values spanned the widest range observed in systems studied here.

**Table 2.**
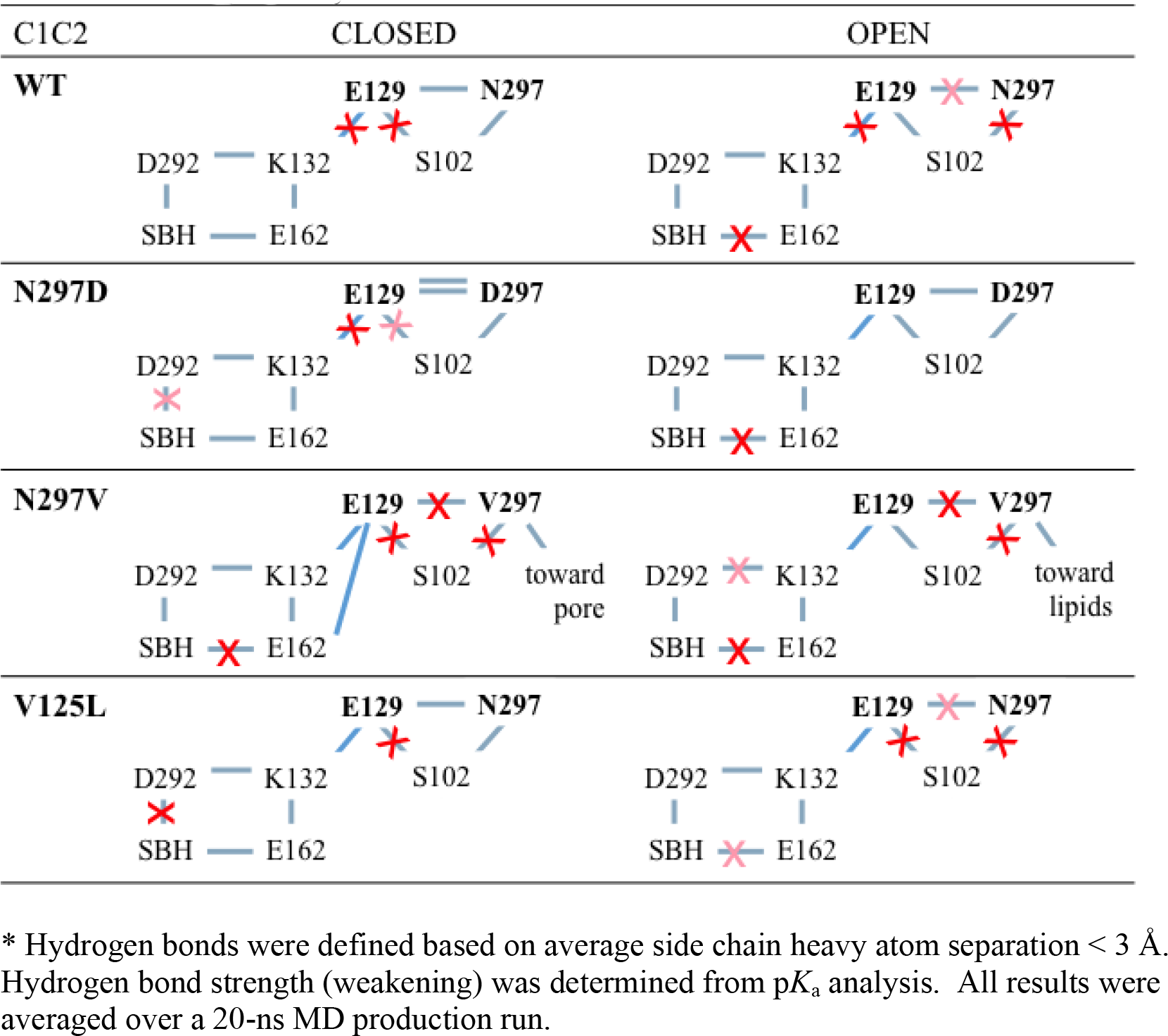
Schematic representation of hydrogen bonds (intact in gray lines, broken in red crosses, weakened in light red crosses) in closed- and open-state C1C2-WT, N297D, N297V, and V125L determined from p*K*_a_ analysis. *

The mean p*K*_a_ value for the gating residue E129 in the closed wildtype C1C2 was mildly basic, p*K*_a_(E129) = 8.8 ± 0.4 (Table S5 and Fig. S2). That value resulted from a single side chain hydrogen bond at the central gate, E129(90)-O…O-N297(258), and weak Coulombic interactions between E129(90) and residues E162(123), D292(253), and K132(93). In contrast, residue E129(90) in the C1C2-N297D mutant was strongly basic, p*K*_a_(E129) = 12.1 ± 0.5, a value more than 3 units higher compared with the wild type channel. That increased p*K*_a_ value resulted from a nearly 4-fold stronger contribution from a double hydrogen bond between side chains of E129(90) and D297(258) in C1C2-N297D compared with a single one in E129-O…O-N297 from the wildtype protein.

Replacement of V125 with leucine resulted in *r*(SB-NH…D292) = 4.0 ± 0.4 Å, an ~1 Å increase in separation between the Schiff base and its proton acceptor when compared with the corresponding distance in N297V closed-state C1C2 variants (Table S4). That structural change led to a 1 p*K*_a_ unit increase in D292 from 2.8 ± 1.2 on average in the wild type to p*K*_a_(D292) = 3.8 ± 1.4 on average in V125L (Table S5). Noting that L125 resides ~10 Å away from D292, it is clear these changes are a consequence of reorganization among intramolecular interactions in C1C2-V125L. In contrast, replacement of polar (asparagine) with electrically charged (aspartic acid) side chain in position 297 (N297D) does not affect the p*K*_a_ of D292 in either closed (p*K*_a_~3) or open (p*K*_a_ ~0.7) states when compared with C1C2-WT (Fig. S2).

#### 3. Structure and hydration of open-state C1C2-WT and mutants

In contrast to the closed-state C1C2 variants with *trans*-retinal, the open-state channels contain retinal in the 13-*cis* conformation. Retinal isomerization caused only small distortions to protein backbone structure (measured as RMSD) and mostly small variations in the separation of gating helices II and VII as reported earlier^25^. Consistent with earlier results, the side chain of E129 (90) had a downward movement when transitioning from the closed to open state. ^14^ Despite the modest structural changes, equilibration of the open-state structures resulted in significant changes to water occupancy (Table S3). For example, waters entered and formed a continuous pathway through each open-state C1C2 (Fig. 1C). While not a complete explanation, specific structural changes occurred upon pore opening that help account for continuous hydration throughout the pore. In general, the number of waters in the transmembrane region increased by an average of 16-27 molecules in each C1C2 variant upon pore opening, bringing the totals up to 53-58 waters. Simultaneously, the maximum distortion to protein backbone structure measured only 0.3 Å (in variant N297D) and the separation between helices II and VII varied by less than ~ 1 Å in most cases.

In the open C1C2-V125L channel, at the intracellular side of the pore, the HII-HVII distance decreased by 0.7 Å. That change marked the only contraction observed. In contrast, the extracellular side expanded by ~ 1 Å. In this variant, the fewest number of waters on average (16) entered the transmembrane region of the pore.

In the open C1C2-N297D mutant, the separation between helices HII-HVII at the extracellular side increased by ~ 1 Å, without contraction elsewhere. That helix expansion was concurrent with a tilt of the R175(136) side chain, and corresponded to water inflow from the extracellular side toward the Schiff base. This mutant attained the highest increase in average water occupancy (+27 H_2_O).

In mutant C1C2-N297V, a 2-Ångstrom-increase occurred in the separation of the gating helices (HII-HVII) at the extracellular side of the pore. Simultaneously, the hydrophobic side chain of residue V297(258) tilted toward the lipid bilayer, allowing space for the influx of an additional 21 waters and formation of a continuous pore.

In the wild-type C1C2, the gating helices expanded by ~ 4 Å at the intracellular side of the pore, representing the largest variation measured. Still, roughly the same number of additional waters entered C1C2-WT (26) as for variant N297D (27). Normal mode analysis identified helix VII as containing the backbone carbon atoms involved in the largest displacements from the equilibrated C1C2-WT structure (Fig. S1). However, our molecular dynamics studies suggest that the change in the separation between helices II and VII does not play a primary role in allowing water access into the previously dry region of the pore. Additional considerations, such as the altered electrostatic charge distribution due to the presence of an acidic residue (D in the N297D mutant), may account for water occupancy even in more restricted areas of the channel, where helices II and VII remain close together.

Based on the C1C2 crystal structure and MD simulations, HVII is the longest of the helices, extending furthest past the lipid bilayer into the intracellular side of the protein. Additionally, retinal is bound to HVII and the putative ion channel is found between helices VII and II. All these factors suggest that HVII would be the most susceptible to conformational changes or displacement during the photocycle, as confirmed by the normal mode analysis (Fig. S1). These changes may occur during transitions between the equilibrated closed and open states studied here. Helix VII is connected to the cytosolic C-terminal of channelrhodopsins (not included in this study), thus it can be expected that even minor conformational changes affecting HVII are likely transduced to the intracellular domain as the first step in a yet to be discovered *C. reinhardtii* signaling pathway. Future simulations of those transitions would help in further defining structural changes that occur during the photocycle.

Results from the MD simulations show that a substantial addition of water molecules accompanies equilibration of the open-state C1C2 structures. While specific structural changes involving expansion/contraction of the separation between gating residues HII and HVII help account for water addition, the comparison is imperfect. For example, the largest expansion between the gating residues of C1C2-WT does not correlate with a similarly larger addition of waters into the transmembrane pore region (Table S3). Noting that rearrangements in protein side chains also accompanied water entry into the open-state channels, we further analyze side chain positions, particularly in the central gating region near residues 129 and 297.

#### 4. Interaction networks in open-state C1C2-WT and mutants

In all open-state C1C2 channels, except N297V, complete pore hydration was achieved by weakening of the closed-state central gate defined by the hydrogen bond between deprotonated E129 and residue 297. The weakened gate, characterized by downward shifts in p*K*_a_ of E129 (Table S5), led to a concurrent increase of hydrophilicity in the gating region. Additional rearrangements in hydrated C1C2 included formation of shorter E129(90)…K132(93)…E162(132) salt bridges in each of the mutants (Table 2). Furthermore, residue D292(253) formed a hydrogen bond with the retinal Schiff base at a close distance of 2.7 ± 0.1 Å (Table S4), while interactions between the Schiff base NH group (SB-NH) and residue E162(123) expanded significantly compared to the closed-state structures. That change identifies D292 as the unambiguous primary acceptor of the Schiff base proton in C1C2.

In contrast to the other C1C2 variants, neither of the gating residues in C1C2-V125L interacted with S102(63) on helix I, following lengthening of the E129…N297 hydrogen bond and pore hydration. Instead, S102(63) formed a hydrogen bond with the backbone oxygen of T98(59) at 2.8 ± 0.3 Å. Concurrently, only in C1C2-V125L did the p*K*_a_(E129) values of both closed (~9) and open (~7) states remain similar to the corresponding values in C1C2-WT.

Mutant C1C2-N297V lacks the central gate in the closed-state and showed a different structure and hydration pattern upon opening, as described above. An alternative formed between E129(90) and E162(123) in the closed state. While a significant Coulombic interaction between E129(90) and E162(123) persisted in the open state (Table S4), the side chain of E129(90) shifted toward S102(63).

In fully hydrated C1C2-N297D, the interaction between E129…D297 weakened as E129(90) formed a salt bridge with K132(93) (Table S4 and double peak in p*K*_a_ (E129, N297D) distribution, red line in Fig. S2)). K132(93) participated in an extended network of salt bridges with E129(90), E162(123), and D292(253), which resulted in p*K*_a_(K132) = 15.4 ± 0.7; an increase of 3 p*K*_a_ units from the corresponding values in closed (11.6 ± 0.3) and open (11.5 ± 0.5) C1C2-WT (Table S5).

Aspartic acid D156 and cysteine C128 were suggested to play an important role in ChR2 kinetics.^26,27^ Interestingly, p*K*_a_ modeling of C1C2 (Figure S2 and Table S5) indicated that the DC gate residues D195(156) and C167(128) were among the amino acids least affected by both single point mutations and pore opening.

## IV. DISCUSSION

Light-activated ion channels, channelrhodopsins (ChRs), are commonly used in optogenetics for selective control of ChR-augmented excitable cells like neurons and muscle cells. While C1C2 has been used as a model of ChR2, recovery of the transient peak current is incomplete and significantly slower for C1C2 when compared with ChR2. To understand better the molecular determinants of channel gating and resulting photocycle kinetics, we conducted computational and electrophysiology experiments on wild type ChR chimera, C1C2-WT, and the following mutants: C1C2-N297D, C1C2-N297V, and C1C2-V125L.

Along with isomerization, retinal deprotonation and reprotonation are key steps in the ChR photocycle. Two residues, E162(123) and D292(253), have been proposed as Schiff base proton acceptors. ^2,8,28^ Subsequently, E123, homologous to D85, the Schiff base proton acceptor in bacteriorhodopsin (bR), was shown to be dispensable for ChR2 function. ^29^ Based on the crystal structure of wild type C1C2, Kato et al. suggested D292(253), analogous to D212 in the bR homologue, as a Schiff base proton acceptor.^17^ In closed-state C1C2-WT equilibrated in our studies, oxygen atoms of either E162(123) or D292(253) resided within ~3 Å of the Schiff base nitrogen. However, in all open-state C1C2 variants, only D292(253) formed a hydrogen bond with the protonated Schiff base, thus confirming D292(253) as a primary Schiff base proton acceptor in C1C2.

The dark-adapted channelrhodopsin central gate, defined by the hydrogen bond between protonated E129(90) and N297(258)^14^ (Table S4), prevents formation of a continuously water-filled channel. In the absence of that interaction, as in the closed state C1C2-N297V mutant, protonated E129 (helix II) forms a hydrogen bond with E162 (helix III) (Table S4), a possible Schiff base proton acceptor in the closed-state structure. That bond creates a barrier to continuous water pore formation in the central region of the pore and acts as an alternative central gate.

Our simulations also show that hydration of the channel requires a minimum of four factors: 1) the presence of 13-*cis* retinal, 2) deprotonation of E129(90), 3) subsequent weakening of the E129(90) and N/D297(258) interaction when present, and 4) water access to previously dry regions among helices I, II, III and VII. When the central gate is absent, as in the N297V variant, then the additional requirement for channel hydration may be associated with an ~2 Å increase in helix II - VII separation at the extracellular side of the pore. This data is consistent with inspection of electrophysiological data that showed that E129 should be deprotonated in the photocycle. ^25^

Opening of the channel occurs via rearrangement of the C1C2 side chains without significant displacement of the protein backbone. Protonation of E129(90) throughout the channelrhodopsin photocycle remains a subject of debate. Recently Hayashi^30^ described an alternative mechanism for C1C2 pore hydration in the presence of protonated E129. Kuhne et al. ^14^ presented an E90-helix II-tilt model that indicates deprotonation of E90 as critical to pore hydration. Here, we observed that the changes in protonation of E129(90) and retinal conformation induce reordering of side chains and an increase in local hydrophilicity, which permit water influx into the transmembrane region among helices I, II, III and VII. These observations agree with the work of Fonfria et al. ChR channel hydration is a prerequisite for transport of monovalent and divalent cations down their electrochemical gradients. Electrophysiology experiments performed here indicate that the N297D mutation in C1C2 slightly increases Na^+^ permeability, while N297V slightly decreases Na^+^ permeability, as expected from changes in the electrostatic interactions between ions and their ligating chemical groups. ^31^

Single point mutations in the gating region of channelrhodopsins strongly influence photocycle kinetics. For instance, replacement of asparagine with aspartic acid in C1C2-N297D yielded overall faster kinetics compared with C1C2-WT. We speculate that aspartic acid in position 297, also in the spatial vicinity of the Schiff base proton acceptor D292(253), contributes to proton transfer. Replacement of asparagine with valine in the analogous position of the ChR2-N258V variant in previous work led to protein expression in oocytes, but negligible photocurrents prevented functional studies of the variant.^16^ In the present work, the introduction of a hydrophobic residue in the gating region of C1C2-N297V resulted in pore gating via E129(90)…E162(123) hydrogen bonding, an alternative central gate. The N297V mutation affected C1C2 photocycle kinetics minimally when compared with wild type C1C2. We attribute the slightly faster off kinetics and recovery of peak current in the dark to weakening of the D292(253)…K132(93) hydrogen bond in the open state compared with other variants studied here. In all studied closed and open mutants, K132(93) forms a network of salt bridges with either E129(90) or E162(123). In ChR2, the analogous lysine K93, along with corresponding residues D253, E90, E213, R123, and H249, have been indicated previously as a component of a proton transfer route to the cytosol.^32^ Further investigations are required to understand that process better in C1C2.

Interestingly, the slowest overall photocycle kinetics occurred in the conservative C1C2-V125L mutant. Moreover, peak currents were completely eradicated upon a second light pulse. Comparable behavior was observed in mutation of the analogous residue, ChR2-V86L. ^16^ In that earlier work, the authors hypothesized that leucine 86 may interfere with helix II displacement and thus affect channel gating. In our simulations, we did not observe significant differences in backbone displacement among C1C2-WT and C1C2-V125L in either closed or open states. In the open state C1C2-V125L channel, however, neither S102…E129 nor S102…N297 hydrogen bonds formed, in contrast to the other variants. These differences suggest that in C1C2, where helices I and II differ from the ones in ChR2, residue S102(63) on helix I plays a role in gating kinetics differences among C1C2-WT and the mutants studied here.

## V. CONCLUSIONS

Our comprehensive electrophysiology studies of C1C2 wild type, C1C2-N297D, C1C2-N297V, and C1C2-V125L show that replacement of a single amino acid has significant influence on photocycle kinetics. Complementary computational studies correlate variations in intramolecular interactions near the C1C2 central gating region to results observed experimentally.

The closed-state C1C2 structure includes all-*trans* retinal, protonated E129(90), and protonated Schiff base (SBH) facing toward the extracellular side of the membrane. The central gate, which holds together helices II and VII (E129(90)…N297(258)), prevents formation of a water-filled channel. Mutations near the gate had minimal effect on structure – in particular, separation between the gating helices remained the same. Similar patterns of partial pore hydration and a water discontinuity near the gating residues were observed for all closed channels.

The opening of the channel and initial hydration of the previously dry gating regions between helices I, II, III, and VII of the channel occurs with 1) the presence of 13-*cis* retinal; 2) deprotonation of a glutamic acid gating residue, E129; and 3) subsequent weakening of the central gate hydrogen bond between the same glutamic acid E129 and asparagine N282 in the central region of the pore. In the studies pursued here, the Schiff base remains protonated and oriented as in the closed state.

Introduction of an acidic side chain in the central gate of C1C2-N297D resulted in slightly faster overall kinetics compared to C1C2-WT, but a similar current decay rate. Also, Na^+^ permeability increased slightly, as expected for addition of stronger electrostatic interactions between the ion and ligating chemical groups. In the open C1C2-N297D, an additional ~27 waters entered the transmembrane region, yielding the second largest backbone RMSD relative to fully hydrated C1C2-WT. At the extracellular side of open C1C2-N297D, the separation between helices II and VII increased by only ~1 Å and induced a tilt of the R175(136) side chain and water inflow from the extracellular side toward the Schiff base and its proton acceptor. In contrast to studies of the closed state structure, the Schiff base proton acceptor in the open state is identified unambiguously here as D292(253).

In the absence of the central gate connecting helices II and VII in the C1C2-N297V mutant, an alternative gate in the central region of the pore formed through a hydrogen bond between E129(90) on helix II and E162(123) on helix III. That alternative gate may help explain why the N297V mutation affected C1C2 photocycle kinetics minimally when compared with wild type C1C2.

In C1C2-V125L, the conservative mutation (valine to leucine) was introduced, which is spatially proximal to the central gate and resulted in the slowest overall photocycle kinetics. Moreover, the peak currents were completely eradicated upon a second light pulse. Based on computational results, S102(63) on helix I may play a role in gating kinetics differences among C1C2-WT and the mutants studied here. Specifically, neither S102…E129 nor S102…N297 hydrogen bonds formed in the C1C2-V125L open-state channel, in contrast to the other variants. Those variants had faster photocycle kinetics and peak currents upon a second light pulse. Instead, V125L disrupts the S102 interaction with gating residues in the open state and thus disrupts ion channel gating.

In summary, replacement of helices I and II in ChR2 with corresponding residues from ChR1, resulted in significantly slower and incomplete recovery of C1C2 transient peak current compared with ChR2. Differences also appeared in the N297 mutants. Thus, kinetic results substantiate that replacement of helices I and II in ChR2 with corresponding residues from ChR1, to make the chimera C1C2, affects the kinetics of channelrhodopsin pore gating significantly, making C1C2 a unique channel. This study correlated fine differentiation of the interaction network in the central region of the pore to changes in photocycle kinetics. Our computational studies confirmed D292(253) as a primary Schiff base proton acceptor in C1C2. Moreover, by using both atomistic computational and electrophysiology studies, we showed how small variations via a single-point mutation near the gating region affect C1C2 photocycle kinetics.

## VI. AUTHOR CONTRIBUTIONS

All authors helped plan the research, analyze the results, and write the manuscript. MRV carried out the molecular simulations. LAP carried out the kinetics measurements.

## VII. ACKNOWLEDGEMENTS

This work was performed, in part, at the Center for Integrated Nanotechnologies, an Office of Science User Facility operated for the U.S. DOE’s Office of Science. The authors have no conflicts of interest.

